# Broadly reactive human monoclonal antibodies targeting the pneumococcal histidine triad protein protect against fatal pneumococcal infection

**DOI:** 10.1101/2021.02.17.431651

**Authors:** Jiachen Huang, Aaron D. Gingerich, Fredejah Royer, Amy V. Paschall, Alma Pena-Briseno, Fikri Y. Avci, Jarrod J. Mousa

## Abstract

*Streptococcus pneumoniae* remains a leading cause of bacterial pneumonia despite the widespread use of vaccines. While vaccines are effective at reducing the incidence of most vaccine-included serotypes, a rise in infection due to non-vaccine serotypes, and moderate efficacy against some vaccine included serotypes have contributed to high disease incidence. Additionally, numerous isolates of *S. pneumoniae* are antibiotic or multi-drug resistant. Several conserved pneumococcal proteins prevalent in the majority of serotypes have been examined as vaccines in preclinical and clinical trials. An additional, yet unexplored tool for disease prevention and treatment is the use of human monoclonal antibodies (mAbs) targeting conserved pneumococcal proteins. Here, we isolate the first human mAbs (PhtD3, PhtD6, PhtD7, PhtD8, PspA16) against the pneumococcal histidine triad protein (PhtD), and the pneumococcal surface protein A (PspA), two conserved and protective antigens. mAbs to PhtD target diverse epitopes on PhtD, and mAb PspA16 targets the N-terminal segment of PspA. The PhtD-specific mAbs bind to multiple serotypes, while mAb PspA16 serotype breadth is limited. mAbs PhtD3 and PhtD8 prolong the survival of mice infected with pneumococcal serotype 3. Furthermore, mAb PhtD3 prolongs the survival of mice in intranasal and intravenous infection models with pneumococcal serotype 4, and in mice infected with pneumococcal serotype 3 when administered 24 hours after pneumococcal infection. All PhtD and PspA mAbs demonstrate opsonophagocytic activity, suggesting a potential mechanism of protection. Our results provide new human mAbs for pneumococcal disease prevention and treatment, and identify epitopes on PhtD and PspA recognized by human B cells.

## Introduction

*Streptococcus pneumoniae* remains a leading cause of infectious morbidity and mortality despite the widespread use of two vaccines for disease prevention (1). The World Health Organization estimates over 1 million deaths occur worldwide each year due to pneumococcal infection (2). Similar to other respiratory pathogens, individuals below the age of 2 and above 65 years of age are more susceptible to invasive pneumococcal disease (3). In addition, there is also an increased frequency and risk of severe infection in individuals with preexisting conditions, including those with diabetes, chronic obstructive pulmonary disease, cardiovascular diseases, and human immunodeficiency virus (4). Although vaccination is widespread in the developed world, pneumococcal infection is responsible for 30% of adult pneumonia and has a mortality rate of 11-40% (5). Furthermore, in regions of the world with high childhood mortality rates, pneumococcal pneumonia is the cause of death for 20-50% of children (6).

*S. pneumoniae* is a common resident of the upper respiratory tract (7), and pneumococcal carriage precedes active infection (8). In young children, carriage rates of *S. pneumoniae* can be as high as 40-60% (9). Colonization is typically asymptomatic, however, *S. pneumoniae* can rapidly disseminate, often following a primary infection such as influenza (10) or COVID-19 (11), to cause pneumonia and invasive disease. Repeated colonization with *S. pneumoniae* typically results in immunization, and several studies have determined that colonization induces serum antibody responses to the capsular polysaccharide (12), and both serum antibody (13–17) and cellular immune responses to protein antigens (18, 19). These antibody levels in serum increase during the first few years of life (16), but tend to decrease in the elderly (20), which may contribute to the higher risk of disease in children and the elderly.

The majority of *S. pneumoniae* isolates are encapsulated, and 100 capsular serotypes have been identified (21), which are based on differences in the chemical structures of the capsular polysaccharide in each serotype (22). Current vaccines are based on eliciting opsonophagocytic antibody responses to the capsular polysaccharide, and utilize either a 13-valent diphtheria toxoid conjugate vaccine to elicit T-dependent, high-affinity, and class- switched antibody responses (PCV13), or a 23-valent capsular polysaccharide mixture (PPSV23) to elicit T-independent antibody responses, or as a booster to PCV13. Anti-glycan antibodies produced in response to the vaccine are serotype-specific due to the distinct chemical structures of the capsular polysaccharides (23). Although vaccines have been highly effective at reducing the incidence of pneumococcal disease, a rise in the incidence of non- vaccine serotypes has occurred, termed serotype replacement (24). In addition, the incidence of invasive disease due to serotypes 3 and 19A have persisted in some reports despite widespread vaccination (25). In terms of treatment, antibiotic resistance among non-vaccine serotypes has risen, and presents challenges in treating pneumococcal infection (26). Based on the limitations of current vaccines and treatments, additional options are currently being explored. For many years, such research has focused on developing vaccines that are broadly reactive, primarily based on the idea that conserved protein antigens present in the majority of pneumococcal serotypes would be effective at preventing disease independent of serotype (27). Multiple antigens have been tested in preclinical infection models, with several entering clinical trials, including the toxin pneumolysin, pneumococcal surface protein A (PspA), pneumococcal surface antigen A (PsaA), pneumococcal choline binding protein A (PcpA), PcsB, serine threonine kinase protein (StkP), and pneumococcal histidine triad protein (PhtD) (28).

PhtD is a member of a group of conserved surface proteins on *S. pneumoniae* that also includes PhtA, PhtB, and PhtC, all of which share histidine triad motifs (29). The proteins have high sequence homology to each other, and PhtB and PhtD share 87% sequence homology (30). PhtD is highly conserved, varying 91-98% among strains isolated from invasive disease cases in children (31). One study of 107 pneumococcal strains showed PhtD was expressed in 100% of tested serotypes (30), while other studies have found PhtD is widely prevalent but is absent in a subset of isolated strains (32–34). The function of the Pht family of proteins has not been fully elucidated, although data has implicated the proteins in attachment of *S. pneumoniae* to respiratory epithelial cells (35, 36). In addition, the first histidine triad motif of PhtD has been shown to be important for zinc acquisition and bacterial homeostasis (37). Although the full structure of PhtD has not been determined, a crystal structure of the third histidine triad motif bound to Zn^2+^, and a solution NMR structure of the N-terminal fragment of PhtD has been determined (38, 39). All Pht proteins are immunogenic and induce protective humoral immunity, and vaccination with these proteins was shown to reduce colonization, sepsis, and pneumonia (29, 40, 41). PhtD has been shown to protect against systemic pneumococcal disease in a mouse model (29), and immunization of rhesus macaques with PhtD along with detoxified pneumolysin protected the animals against pneumococcal infection (42). Fragments of PhtD have also been assessed for protective efficacy, and somewhat conflicting reports have demonstrated that both the N and C terminus are immunogenic and protective (43, 44). PhtD was recently used as an antigen in a phase IIb clinical trial, demonstrating that PhtD remains an antigen of interest in pneumococcal vaccinology, although PhtD was administered along with PCV13, so a direct comparison of PhtD vs PCV13 was not accomplished (45). Mouse monoclonal antibodies to PhtD were shown to protect mice using a macrophage and complement dependent mechanism (46), and human polyclonal antibodies to PhtD were shown to reduce adherence of the pneumococcus to lung epithelial cells and reduce murine nasopharyngeal colonization (47). Human polyclonal antibodies generated in response to alum adjuvanted PhtD vaccination were also shown to protect mice from pneumococcal disease (48).

Another vaccine antigen, PspA, is an important virulence factor of *S. pneumoniae* and one of the most abundant surface proteins (49). As with PhtD, PspA is found in the majority of examined clinical isolates (33, 50). PspA mutant strains are cleared faster from the blood of mice compared to intact strains (51), and vaccination with PspA protects mice from pneumococcal infection (52–58). PspA is less conserved than PhtD, and is grouped into three families with >55% identity, and six clades with >75% identity (59). PspA has four distinct structural domains, including the alpha-helical region, the proline rich region, the choline binding repeat domain, and the cytoplasmic tail, of which the proline rich region is highly conserved across clades, while the N-terminal alpha-helical region is more variable (60). PspA has been shown to inhibit complement deposition (61–63), and has shown specificity for binding of human lactoferrin, although the importance of this binding is unclear (64). An X-ray crystal structure of the lactoferrin binding domain of PspA in complex with the N-terminal region of human lactoferrin has been determined (65). Mouse mAbs to PspA have been shown to prolong survival of mice, and improve efficacy of antibiotic treatment (64). Additionally, antibodies isolated from humans following immunization with recombinant PspA are broadly cross-reactive and protect mice from pneumococcal infection with heterologous PspA (66, 67). A clinical trial of a recombinant attenuated salmonella typhi vaccine vector producing PspA has been completed (NCT01033409), and a protein-based Phase Ia clinical trial incorporating PspA is current underway (NCT04087460).

It is well-defined that antibodies can prevent pneumococcal infection based on the success of antibody-based pneumococcal vaccines. Since both PspA and PhtD are protective antigens, and elicit protective antibodies, it is reasonable to assume that human mAbs to these antigens would be protective. As these proteins are highly conserved across pneumococcal serotypes, mAbs to PhtD and PspA could prevent and possibly treat disease from a broad-spectrum of pneumococcal serotypes. Human mAbs are promising as therapeutics for bacterial pathogens, as bezlotoxumab was FDA approved for prevention of recurrent *Clostridium difficile* infection (68). However, there have been no human mAbs isolated to any pneumococcal protein antigens. Serum antibodies to PhtD and PspA are elicited in response to pneumococcal carriage (16, 69, 70), and in this study, we generated human monoclonal antibodies to PhtD and PspA from healthy human subjects. We determined the serotype breadth and epitope specificity of the mAbs, and demonstrated the protective efficacy of PhtD-specific human mAb in multiple mouse models of pneumococcal infection.

## Results

### Isolation of pneumococcal protein-specific human mAbs

To identify PhtD and PspA-specific human mAbs, we recombinantly expressed His-tagged PhtD and PspA from strain TCH8431 (serotype 19A) in *E. coli* **(Figure 1A)**, and utilized these proteins to screen stimulated B cells from human donor peripheral blood mononuclear cells (PBMCs) as previously described (71). PBMCs from healthy human subjects were plated onto a feeder layer expressing human CD40L, human IL-21, and human BAFF for six days to stimulate B cell growth and antibody secretion. Cell supernatants from the stimulated B cells were screened against recombinant PhtD and PspA by enzyme-linked immunosorbent assay. Responses to the recombinant proteins were varied between subjects as shown in an example in **Figure 1B**. From five subject PBMCs, we fused several reactive wells for generation of human hybridomas and subsequent human mAb isolation. Four hybridomas lines, each from unique donors, were successfully generated and biologically cloned by single cell sorting for PhtD, and one mAb was generated for PspA from an independent subject. Each of the mAbs to PhtD had similar binding EC_50_ values determined by ELISA **(Figure 1C)**, and mAbs to PhtD and PspA bound with high avidity with EC_50_ values ranging from 26-45 ng/mL **(Figure 1C, 1D)**. To determine the V, D, and J genes utilized by each mAb, the hybridomas were sequenced by RT-PCR followed by TA cloning and the results are shown in **Table 1**, and the sequences are provided in supplemental material. mAbs PhtD3 and PhtD6 utilize kappa light chains, while mAbs PhtD7 and PhtD8 use lambda light chains. All mAbs were of the IgG_1_ isotype based on isotyping data determined by ELISA. All mAbs utilize unique heavy chain and light chain V genes, with the exception of mAbs PhtD7 and PhtD8, as these share predicted V_L_ and J_L_ gene usage, although LCDR3 sequences share little sequence identity. mAbs PhtD7 and PhtD8 share V_H_ and J_H_ gene usage, although vary in the use of the D_H_ gene, which leads to stark differences in CDR3 lengths, with mAbs PhtD7 and PhtD8 having 20 amino acid and 8 amino acid length HCDR3 lengths, respectively. mAb PspA16 shares V gene usage with mAbs PhtD7 and PhtD8.

**Table 1:** Summary of genetic characteristics of pneumococcal-specific mAbs.

|  |  |  |  |  |  |  |  |  |
| --- | --- | --- | --- | --- | --- | --- | --- | --- |
| mAb | isotype | V<sub>H</sub> gene<br>(% mutation) | D<sub>H</sub> | J<sub>H</sub> | HCDR3<br>sequence | V<sub>L</sub><br>(% mutation) | J<sub>L</sub> | LCDR3<br>sequence |
| **PhtD3** | IgG<sub>1</sub>, κ | V1-69*18<br>(85%) | D3-16*01 | J3*01 | ARDGHIMRTTLSDAALDV | V3-20*01 (91%) | J4*01 | QQYQNSPFT |
| **PhtD6** | IgG<sub>1</sub>, κ | V1-8*01<br>(93%) | D2-15*01 | J5*02 | ARGPYWVENWFDT | V1-39*01 (92%) | J1*01 | QQSYSNQKT |
| **PhtD7** | IgG1, λ | V1-2*02<br>(91%) | D3-16*01 | J4*02 | ARVLRGSYDFRGNYPHDFDY | V4-69*01 (88%) | J3*02 | QTWDTGLQGV |
| **PhtD8** | IgG<sub>1</sub>, λ | V1-2*02<br>(93%) | D2-15*01 | J4*02 | ARGGTLDH | V4-69*01 (94%) | J3*02 | HTWVTNIHLV |
| **PspA16** | IgG<sub>1</sub>, κ | V1-2*02<br>(93%) | D3-10*01 | J1*01 | ARAWAPGAEYLHH | V3-20*01 (94%) | J3*01 | QQHDHSPFT |
| Analysis was performed using IMGT/V-Quest. |  |  |  |  |  |  |  |  |

**Figure 1.**
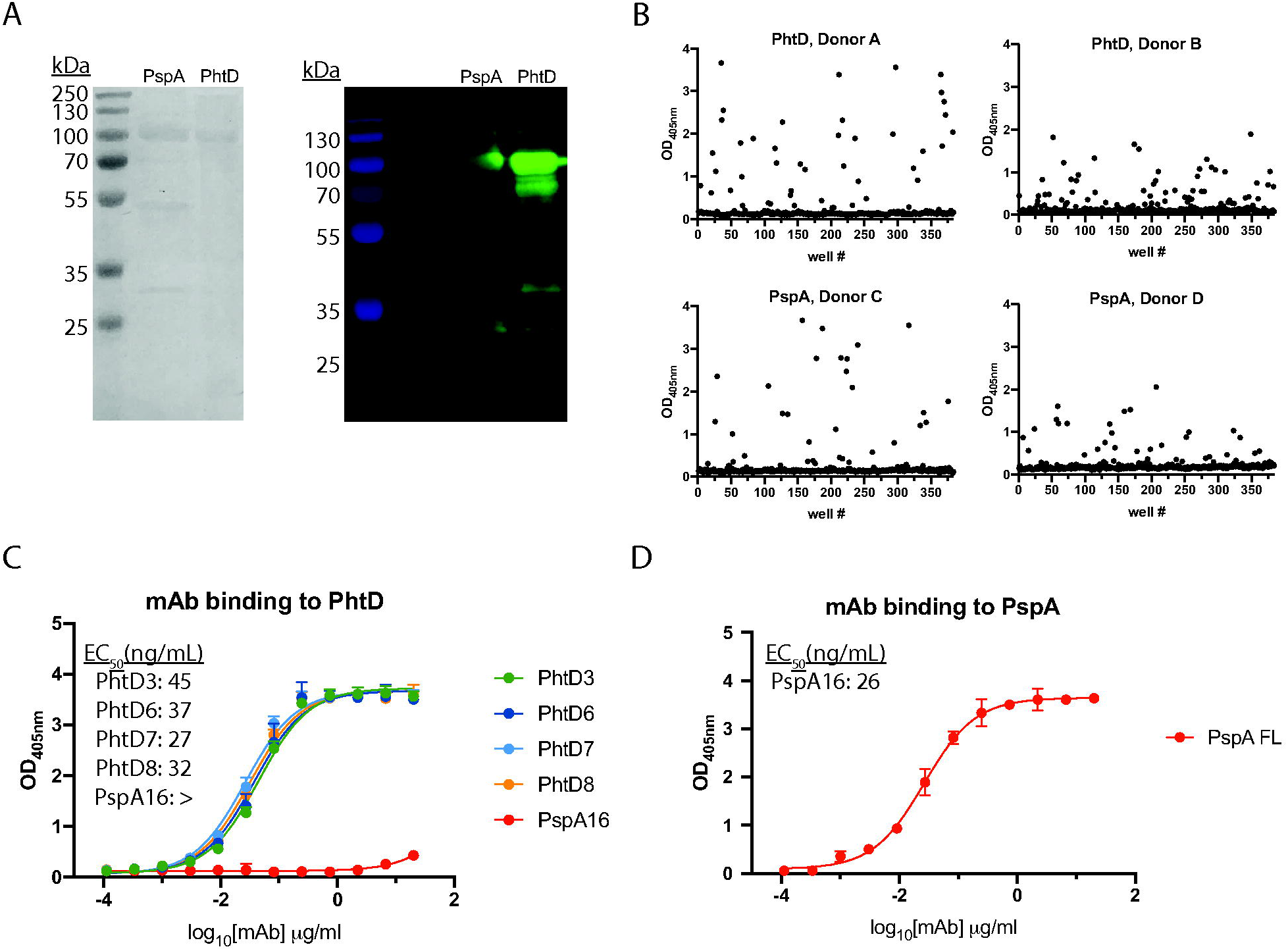
Antibody responses and mAb binding properties to recombinant PhtD and PspA proteins. (A) SDS-PAGE (left) and western blot (right) of purified recombinantly expressed PspA and PhtD. Both proteins were pure with the appearance of degradation products. (B) ELISA binding responses from the supernatant of stimulated B cells to recombinant PhtD and PspA proteins. (C) ELISA binding curves of anti-PhtD mAbs against recombinant PhtD protein. PspA16 was utilized as a negative control. > indicates no binding was observed at an OD_405_ over 1 Abs at the highest concentration. (D) Binding of PspA16 to recombinant PspA. For (C) and (D), computed EC_50_ values in ng/mL are reported from a non- linear regression curve fit (agonist). Data points indicate the average of four replicates from one of at least two independent experiments. Error bars indicate 95% confidence intervals. Summary of genetic characteristics of pneumococcal-specific mAbs.

### Epitope mapping of the human mAbs

To identify the specific regions of PhtD targeted by the human mAbs, we generated truncated fragments of PhtD based on a secondary structure predictor. The fragments were fused to the maltose binding protein (MBP) to ensure solubility, and expressed in *E. coli* and purified using amylose resin. The majority of the fragments were >90% pure with the exception of free MBP protein for the MBP fusion proteins **(Figure 2A)**. To identify the specific regions of PhtD targeted by the isolated mAbs, we measured ELISA binding of mAbs to fragments of PhtD. Since there are no previous mAbs that have been generated to these proteins with defined epitopes, the generated fragments provide rough estimates of mAb epitopes. Each of the four mAbs bind to a unique region on the PhtD protein **(Figure 2B)**. mAbs PhtD3 and PhtD6 bind the N-terminal portion of the protein, while mAb PhtD8 binds the C-terminal portion. mAb PhtD7 appears to target a unique conformational epitope that is dependent on amino acids 341- 838, but this mAb does not bind 341-647 or 645-838 fragments. We next assessed the epitopes of the mAbs by competitive biolayer interferometry to compare the binding epitopes between mAbs. Anti penta-His biosensors were loaded with His-tagged PhtD protein, and mAbs were competed for binding sequentially **(Figure 2C,2D)**. The mAbs bind distinct regions with limited competition similar to results from the fragment ELISA data. mAbs PhtD3 and PhtD6 show intermediate competition, and the epitopes for these mAbs also overlap in our fragment ELISA data. To map the binding region of mAb PspA16, we fragmented PspA into several truncations based on previously determined domains **(Figure 3A)** (60). mAb PspA16 had high avidity to recombinant PspA, and bound to the N-terminal fragment 1-247 based on positive binding to amino acid fragments 1-438 and 1-512 and negative binding to 247-512, 436-725, and 247-725 fragments **(Figure 3B)**.

**Figure 2.**
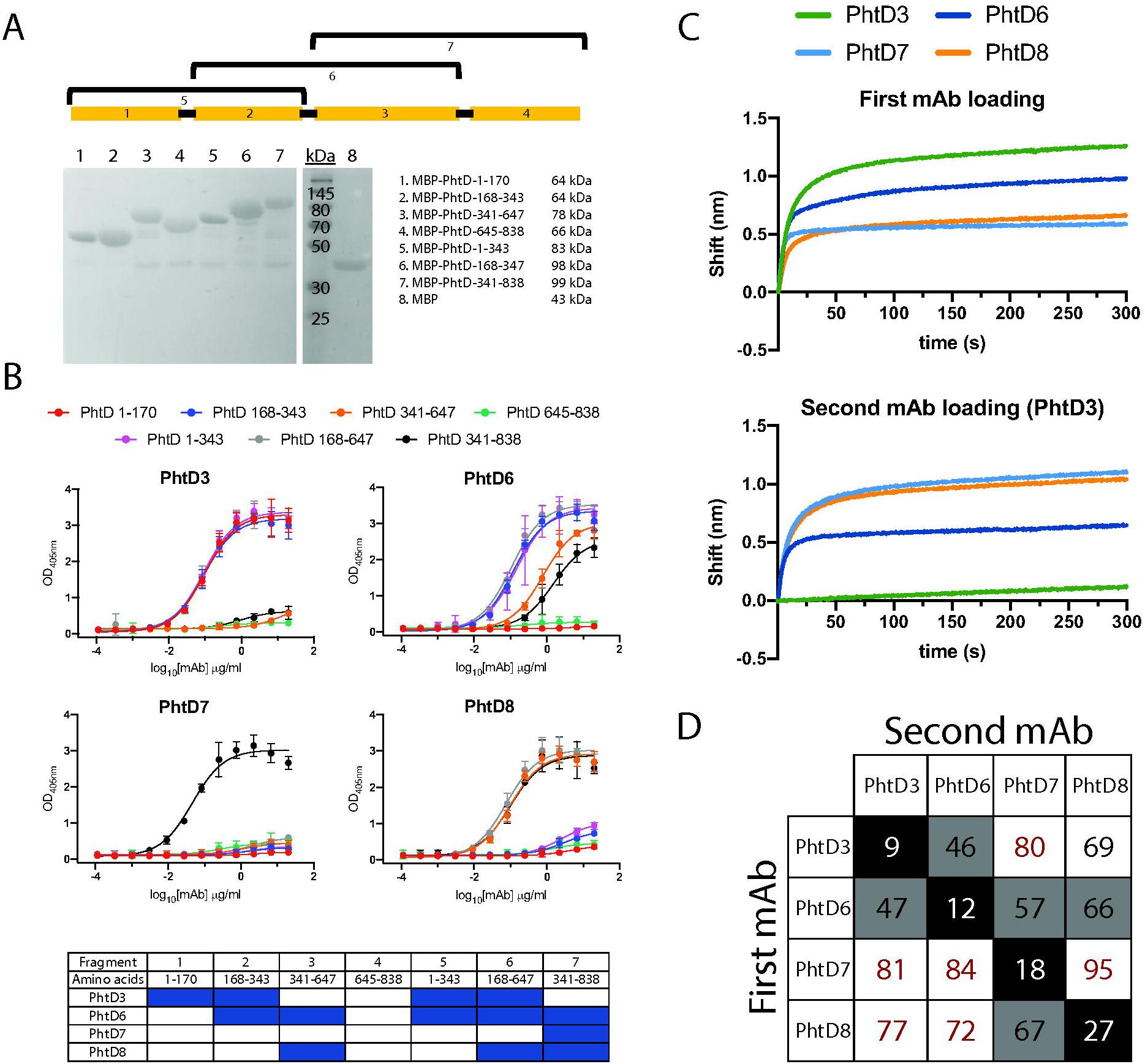
Epitope mapping of anti-PhtD mAbs. (A) SDS-PAGE of the purified maltose binding protein (MBP) PhtD fragment fusion proteins. Each fusion protein was pure after purification with the exception of free MBP. (B) ELISA binding curves of the PhtD mAbs to each MBP-PhtD fragment. A summary of the binding curves is displayed below the binding curves, where cells colored in blue indicate binding, and those colored in white indicate no binding. Data points indicate the average of four replicates from one of at least two independent experiments. Error bars indicate 95% confidence intervals. (C) An example of an epitope mapping experiment for the anti-PhtD mAbs. The top graph displays the signal from biolayer interferometry of the first mAb loaded onto immobilized PhtD protein. The signal for each mAb is colored according to the legend. The bottom graph displays the signal from loading of the second mAb in the presence of the first mAb, mAb PhtD3 in this example. A decrease in signal compared to the top graph is observed for mAbs PhtD3 and PhtD8, as mAb PhtD3 competes with itself, and mAbs PhtD3 and PhtD6 have partially overlapping epitopes. (D) Epitope mapping of the PhtD-specific mAbs. Data indicate the percent binding of the competing antibody in the presence of the primary antibody, compared with the competing antibody alone. Cells filled in black indicate full competition, in which ≤33% of the uncompeted signal was observed; cells in gray indicate intermediate competition, in which the signal was between 33% and 66%; and cells in white indicate noncompetition, where the signal was ≥66%.

**Figure 3.**
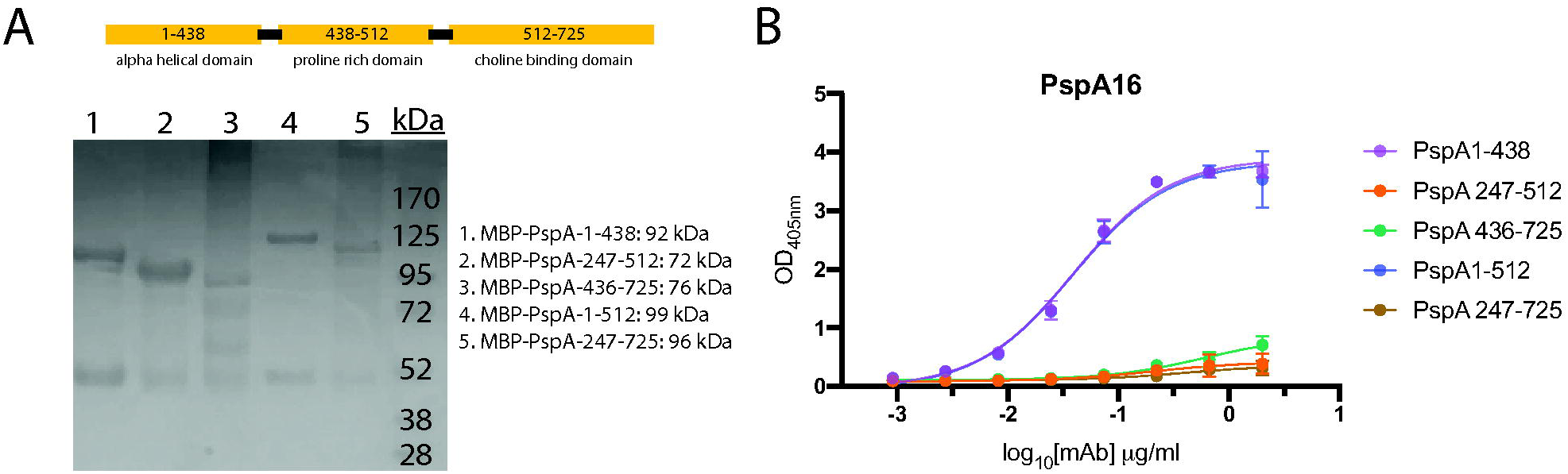
Epitope mapping of PspA16. (A) SDS-PAGE of recombinant MBP PspA fragment fusion proteins. Fragments 1-2 and 4-5 purified well, with only visible MBP protein as a contaminant. PspA fragment 3 has multiple co-purified bands and/or degradation products. (B) ELISA binding curves for PspA16 to each fragment. PspA16 bound to fragment 1 and fragment 4, but not others, suggesting the epitope lies within amino acids 1-247. Data points indicate the average of four replicates from one of at least two independent experiments. Error bars indicate 95% confidence intervals.

### Serotype breadth of the isolated PhtD-specific mAbs

Pneumococcal surface proteins PhtD and PspA are conserved across serotypes, and are widely prevalent in the majority of serotypes. As such, human mAbs to these antigens could have the potential to treat pneumococcal infection from multiple serotypes. In order to determine the serotype breadth of the isolated mAbs, we initially assessed mAb binding to two diverse pneumococcal serotypes, strain TCH8431 (serotype 19A), from which the genes for recombinant PhtD and PspA proteins were cloned and expressed, and the commonly used laboratory strain TIGR4 (serotype 4). PspA shares 88% amino acid sequence identity between TCH8431 and TIGR4, although significant variability is present in the N-terminal domain, with 70% identity in amino acids 1-247. In contrast, PhtD shares 98% amino acid sequence identity between these two strains. We conducted western blots by probing bacterial lysates from TIGR4 and TCH8431 with mAbs PhtD3 and PspA16. mAb PspA16 only labels PspA protein from strain TCH8431 **(Figure 4A)**, while mAb PhtD3 is able to label PhtD protein from both pneumococcal strains. However, as the bacterial lysis likely results in protein denaturation, it is possible the epitope for mAb PspA16 is altered during denaturation. We next determined if mAbs isolated against each of the recombinant proteins bind whole bacteria. We conducted ELISA assays by coating plates with fixed bacteria and measuring mAb binding by ELISA. mAbs PhtD3, PhtD6, PhtD7, and PhtD8 were broadly reactive across multiple unrelated pneumococcal serotypes, and mAbs PhtD3, PhtD6, and PhtD7 had higher avidity to fixed bacteria as compared to PhtD8 **(Figure 4B)**. In contrast, PspA16 bound only to strain TCH8431, similar to results from the western blot experiments. Since PspA16 binds to the most variable region of PspA, the reduced binding to divergent serotypes was expected. In a third experiment, we assessed binding of the mAbs to a panel of pneumococcal serotypes by flow cytometry. As shown in **Figure 4C**, we utilized serum from a donor vaccinated 21 days previously with Prevnar-13 as a positive control. The PhtD mAbs bound to the majority of tested serotypes, with mAbs PhtD3 and PhtD8 showing the broadest binding. In contrast, PspA16 bound only to TCH8431 and the serotype 3 strain WU2.

**Figure 4.**
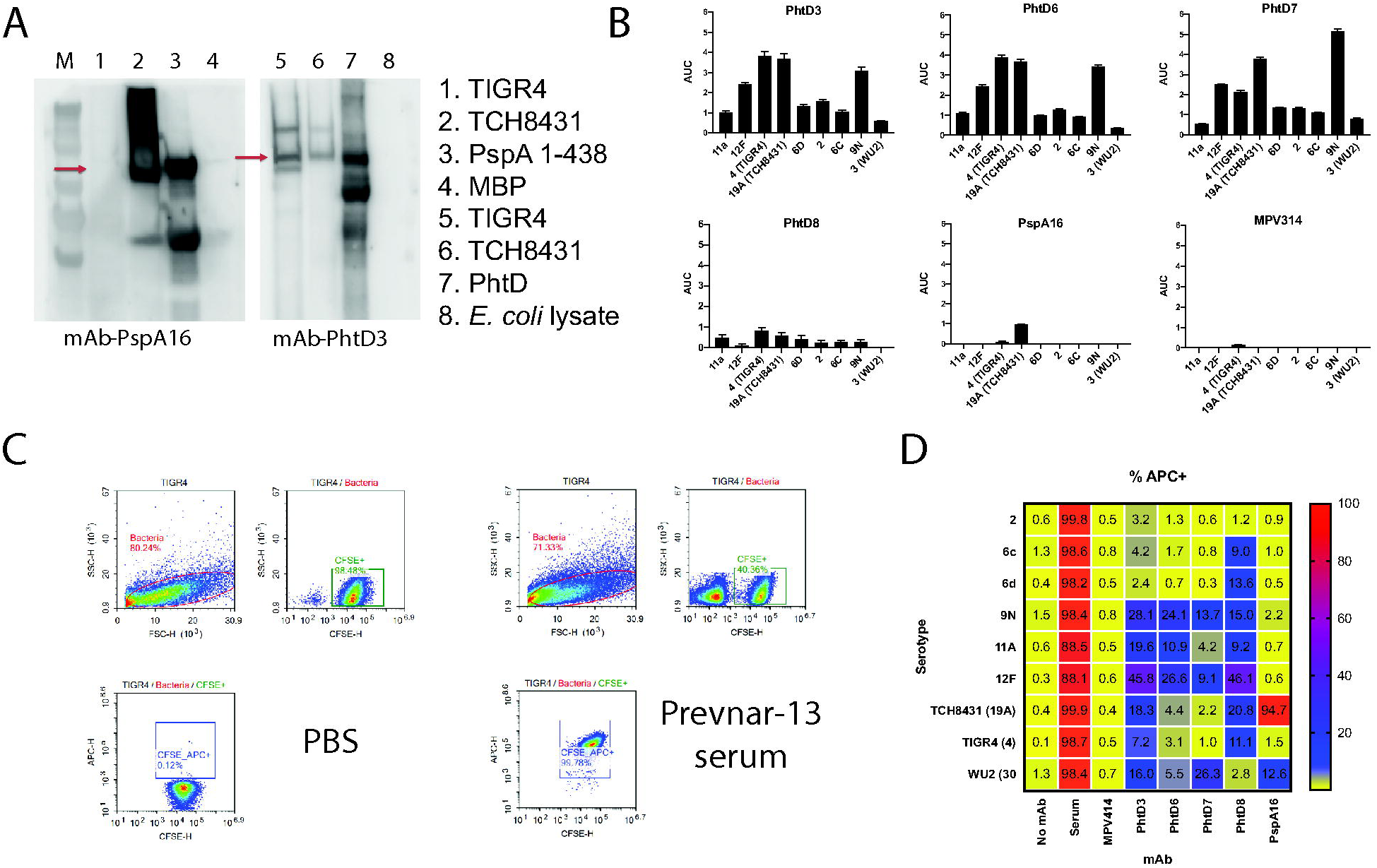
Serotype breadth of the isolated mAbs. (A) Western blot of TIGR4 and TCH8431 strains with PspA16 and PhtD3 as the primary antibodies. In the western blot for PspA16, PspA fragment 1-438 fused to the MBP was used as the positive control, and MBP was used as the negative control. In the PhtD3 western blot, recombinant PhtD was used as the positive control, and *E. coli* lysates were used as the negative control. (B) Area under the curve (AUC) values calculated from ELISA binding curves of serially diluted mAbs against plates coated with fixed bacteria. The ELISA binding curves were the the average of four data points from one of at least two independent experiments. The baseline for the AUC calculation was set as the average of the signal for the highest concentration (20 µg/mL) of the negative control mAb MPV314. Error bars are the standard error from the AUC calculation. (C) Example gating strategy for antibody binding to bacteria. Bacteria were labeled with CFSE, and antibodies were labeled with APC. (D) Heat map and percentages for antibody binding to each pneumococcal serotype. Data are averages from 3-4 experiments, and are the percent of bacteria that are APC-positive. MPV314 and MPV414 are human antibodies specific to the human metapneumovirus fusion protein, and these were used as negative controls.

### PhtD3 protects mice from fatal pneumococcal infection

As mAbs PhtD3 and PhtD8 exhibited the highest overall breadth in the serotype binding analysis by flow cytometry, these mAbs were further analyzed for protective efficacy in the mouse model. In addition, these mAbs were chosen in order to identify if the epitope specificity of mAbs to PhtD affect protective efficacy, as they target nonoverlapping epitopes. Mouse mAbs to PhtD (46) and polyclonal human antibodies from both healthy human subjects (47) and PhtD- vaccinated humans (48) have been shown to protect against colonization or disease in mouse models of pneumococcal infection. However, no human mAbs have been examined for protective efficacy. To determine if the PhtD-specific mAbs protect against infection, we examined the efficacy of mAbs PhtD3 and PhtD8 in a mouse model of pneumococcal pneumonia with a serotype 3 strain (WU2), as serotype 3 is a leading cause of invasive pneumococcal disease (72). Since the mAbs were isolated from human hybridomas, and thus have authentic human Fc regions, we isotype-switched the Fc region to the closest mouse homolog (human IgG_1_ became mouse IgG_2a_). mAbs PhtD3 and PhtD8 chimeras with mouse IgG_2a_ Fc regions (PhtD3-IgG_2a_ and PhtD8-IgG_2a_) were recombinantly expressed in HEK293F cells for testing in the mouse model. As a control for the study, we purchased a mouse IgG_2a_ isotype control antibody. We first examined the binding of the mAbs to ensure binding was still observed for the recombinant PhtD3-IgG_2a_ and PhtD8-IgG_2a_ mAbs, and that no binding was observed for the isotype control mAb. As expected, PhtD3-IgG_2a_ and PhtD8-IgG_2a_ had similar binding avidity to recombinant PhtD as hybridoma-derived PhtD, while the isotype control showed no binding **(Figure 5A)**. We first tested the prophylactic efficacy of PhtD3-IgG_2a_ and PhtD8-IgG_2a_ in a pneumonia model with pneumococcal serotype 3. Both mAbs prolonged the survival of mice compared to the isotype control, although those mice treated with mAb PhtD3 demonstrated higher survival (80% versus 30%) **(Figure 5B, 5C).** As mAb PhtD3-IgG_2a_ protected a larger percentage of mice, we chose this mAb for further analysis. mAb PhtD3-IgG_2a_ was then tested for protective efficacy against pneumococcal serotype 4 (TIGR4) to identify if the broad binding correlates to broad protection. In experiments with TIGR4, we used only an isotype control mAb group since no significant difference was observed between the PBS and isotype control mAb groups in the serotype 3 experiments. For this serotype, we used CBA/N mice for the intranasal infection model as TIGR4 was not sufficiently lethal by intranasal infection in C57BL/6 mice. CBA/N mice have previously been shown to be susceptible to serotype 4 (73). PhtD3-IgG_2a_ prolonged survival of mice, providing 93% protection compared to 47% for the isotype control **(Figure 5C)**. As we were not able to test PhtD3-IgG_2a_ in an intranasal infection model with TIGR4 in C57BL/6 mice, we conducted an experiment in C57BL/6 mice in which mice were intravenously infected with TIGR4 to model septic pneumococcal infection. In this study, PhtD3-IgG_2a_ prolonged survival of mice with 69% efficacy compared to 27% survival with the isotype control **(Figure 5D)**. The most clinically relevant scenario for mAb treatment of pneumococcal infection would be administration after pneumococcal infection. To model such a scenario, we infected mice with pneumococcal serotype 3, and administered mAb PhtD3-IgG_2a_ 24 hrs after infection. In this model, 65% of PhtD3-IgG_2a_ treated mice survived the infection compared to 10% for the isotype control group **(Figure 5E).**

**Figure 5.**
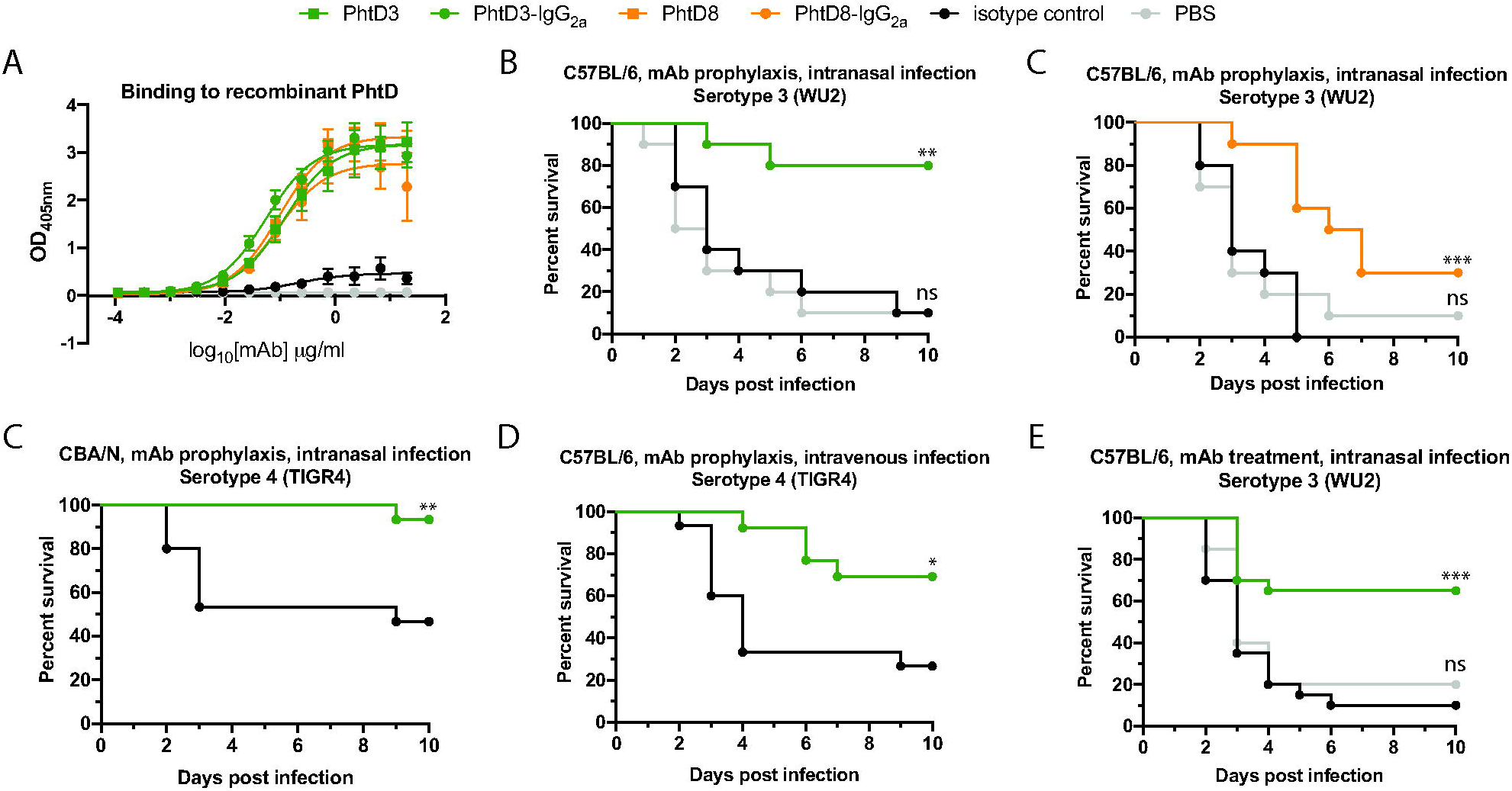
Protective efficacy of anti-PhtD mAbs. (A) ELISA binding curve of mAb PhtD3, the isotype-switched mAb PhtD-IgG_2a_, and an IgG_2a_ isotype control. Data points indicate the average of four replicates from one of at least two independent experiments. Error bars indicate 95% confidence intervals. (B) Prophylactic efficacy of mAb PhtD3 in an intranasal infection model of pneumococcal serotype 3 (strain WU2) in C57BL/6 mice. **, P=0.0012, ns=not significant via log-rank (Mantel-Cox) test. n=10 mice/group. (C) Prophylactic efficacy of mAb PhtD8 in an intranasal infection model of pneumococcal serotype 3 (strain WU2) in C57BL/6 mice. ***, P=0.0009, ns=not significant via log-rank (Mantel-Cox) test. n=10 mice/group. (C) Prophylactic efficacy of mAb PhtD3 in an intranasal infection model of pneumococcal serotype 4 (strain TIGR4) in CBA/N mice. **, P=0.0045 via log-rank (Mantel-Cox) test. n=15 mice/group. (D) Prophylactic efficacy of mAb PhtD3 in an intravenous infection model of pneumococcal serotype 4 (strain TIGR4) in C57BL/6 mice. **P=0.0101 via log-rank (Mantel-Cox) test. n=13-15 mice/group. (E) Treatment efficacy of mAb PhtD3 in an intranasal infection model of pneumococcal serotype 3 (strain WU2) in C57BL/6 mice. ***P=0.0002, ns=not significant via log-rank (Mantel-Cox) test. n=20 mice/group.

### PhtD-specific human mAbs have opsonophagocytic activity

The correlate of protection for current pneumococcal vaccines is based on the elicitation of anti-capsule antibodies that opsonize bacteria, leading to their phagocytosis by host immune cells and subsequent bacterial killing (74, 75). Mouse mAbs isolated by vaccination with PhtD were previously shown to induce bacterial opsonophagocytosis, which was dependent on complement and macrophages (46). To determine a potential mechanism of protection by PhtD3, and additional PhtD mAbs, we utilized established opsonophagocytosis killing assays (OPKAs) using the HL-60 cell line. We tested the mAbs against serotypes 4 (strain TIGR4), 3 (strain WU2), and serotype 19A (strain TCH8431), from which our PhtD and PspA constructs were cloned. These mAbs were also compared to purified IgG obtained from a human subject previously vaccinated with Prevnar-13 21 days before blood collection, as the OPKA assay is the standard to measure vaccine uptake (76). All PhtD mAbs induced decreased colony forming units against all three serotypes compared to no antibody and an irrelevant mAb to human metapneumovirus **(Figure 6A)**. PspA16 also decreased colony forming units against all three serotypes, although the efficacy against serotype 4 was lower as expected based on the serotype binding data. To confirm these findings, we adopted a flow-based assay previously shown to work for group B Streptococcus (77). HL-60 cells were incubated with opsonized bacteria that were labeled with pHRodo, which leads to fluorescent HL-60 cells upon phagocytosis of labeled bacteria. Similar to our results from the OPKA assay, all PhtD mAbs induced an increase in pHRodo+ HL-60 cells compared to no antibody and isotype control antibody analyses **(Figure 6B)**. Purified IgG from an unvaccinated donor showed the highest number of pHRodo+ cells, as human IgG contains antibodies to multiple pneumococcal surface proteins. Interestingly, PspA16 also induced increased uptake to all three serotypes in this assay, although the highest activity was observed for serotype 19A, the serotype from which we cloned our PspA gene.

**Figure 6.**
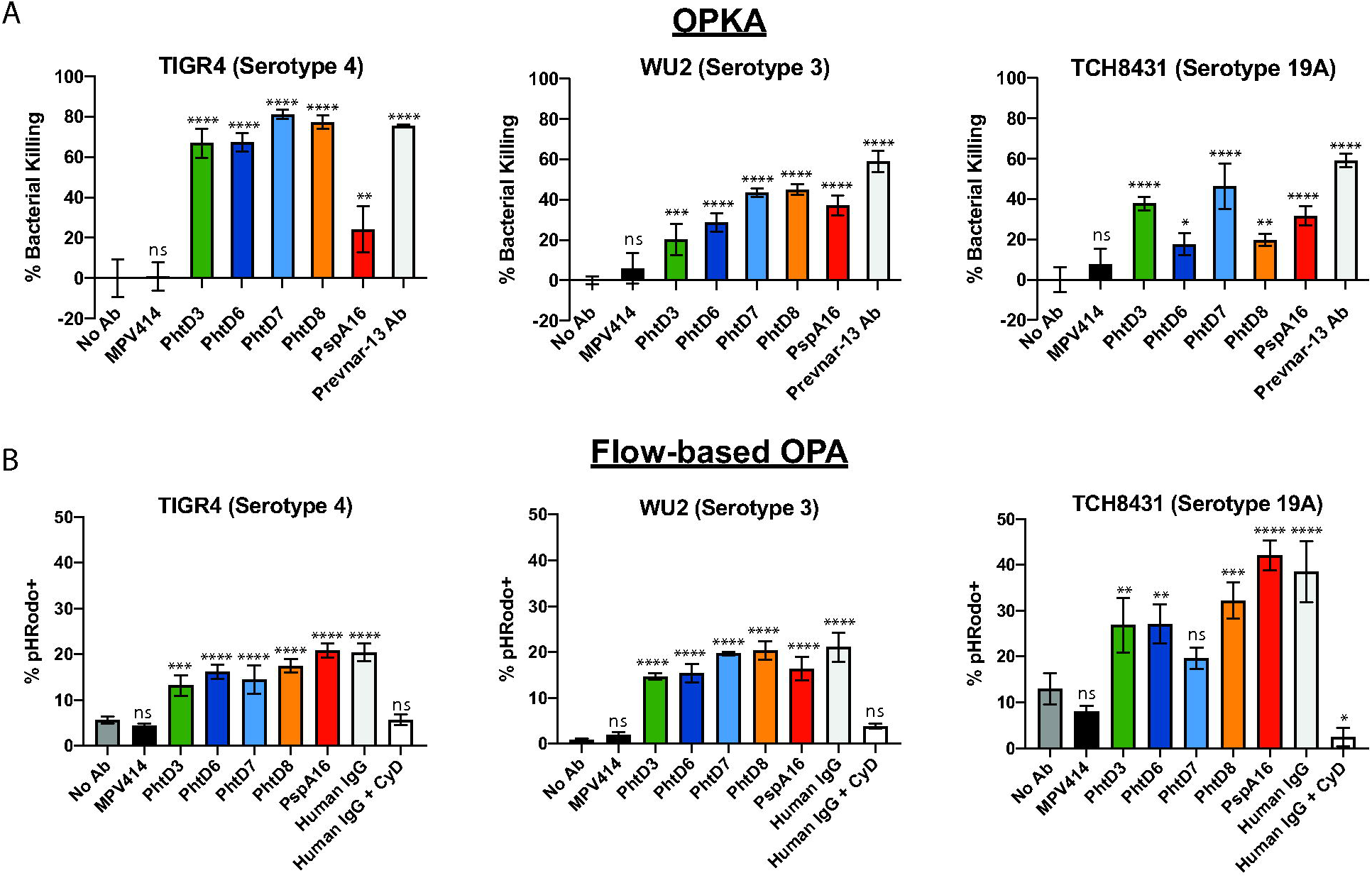
Opsonophagocytic activity of PhtD-specific human mAbs. (A) mAbs and serum were tested in a standard OPA assay using differentiated HL-60 cells. Bacteria were opsonized with antibodies, and subsequently HL-60 cells were added before plating onto blood agar plates. Plates were incubated overnight and CFUs counted. Data are averages of three replicates from one experiment. Error bars represent the range. % Bacterial Killing was calculated as the counted CFU value of each triplicate normalized against the average of the No Ab control. One-way ANOVA analysis with Dunnett’s multiple comparisons test was used to determine significance. ns=not significant, ****P<0.0001. (B) mAbs and serum were tested in a flow-based opsonophagocytosis assay. pHRodo-labeled bacteria were opsonized with antibodies, and incubated with HL-60 cells before being subjected to analysis by flow cytometry. Data indicate the percent of CD38+CD11b+ HL-60 cells that are pHRodo+. Each bar graph is the average of three experimental replicates and error bars are the standard deviation. ns=not significant, ***P=0.0001-0.0006, ****P<0.0001 via one-way ANOVA analysis with Dunnett’s multiple comparisons test. MPV414 is a human mAb specific to the human metapneumovirus fusion protein.

## Discussion

In this study, we have isolated and determined the binding affinity, epitope specificity, serotype breadth, and protective properties of the first human mAbs to any pneumococcal surface protein. Both PhtD and PspA have been examined in depth as vaccine candidates for prevention of pneumococcal infection, although the current outlook for progress of these antigens in the era of conjugate vaccines remains uncertain. However, human mAbs to these conserved antigens offer the ability for pan-serotype recognition and potentially disease prevention and treatment. In contrast, human mAbs isolated following vaccination with pneumococcal polysaccharide vaccines are highly serotype specific (23), and would offer limited use in the clinic due to their highly specific serotype specificity.

Based on the B cell stimulation and screening results, it is clear that healthy individuals have circulating B cells specific to pneumococcal antigens PhtD and PspA. One drawback of our study is the lack of knowledge on the infection history of the human subjects used in the study. It has been previously shown that colonization by *S. pneumoniae* results in immunization, and it is unknown whether all of these donors were previously infected with *S. pneumoniae*. Therefore, the mAbs isolated here are likely the result of pneumococcal colonization, which resulted in class switched B cells with 85-94% somatic mutation, which is similar to previous work in our lab studying healthy individuals who were presumed to be previously infected with human metapneumovirus (71, 78). Each of the PhtD-specific mAbs was isolated from unique human subjects, and mAbs PhtD3, PhtD6, and PhtD7/8 utilized different heavy and light chain V genes. Interestingly, mAbs PhtD7 and PhtD8 utilize the same V gene in both heavy and light chain, yet differ in the predicted heavy chain J gene, which leads to highly different CDR3 lengths. Although these two mAbs share common heavy and light chain V genes, the epitopes for these mAbs do not compete and have only partial overlap based on the binding experiments with truncated protein fragments. It is a striking observation that the N-terminal specificity of mAbs PhtD3 and PhtD6 correlates with higher binding to whole cell bacteria compared to PhtD8, as the N-terminal region of the protein is predicted to be attached to the bacterial surface, leaving the C-terminal half more surface exposed (35). Further mapping experiments through X-ray crystallographic analysis will help clarify this observation. Previous work has identified specific linear peptide epitopes that are immunodominant in pediatric patients with invasive pneumococcal disease, and these included AA 88-107, AA 172-191, and AA 200-219 (79). These peptides overlap with the epitopes for mAbs PhtD3 and PhtD6, although we have not yet determined if these mAbs bind these peptide epitopes. Overall, these data suggest the human antibody response to PhtD targets multiple epitopes. For PspA16, the mAb targets the N-terminal region of PspA, which has a high number of negatively charged residues, and has been shown to be protective in several studies (80). Mouse mAbs isolated against PspA were determined to target the N-terminal fragment as well, suggesting this domain is immunogenic in both mice and humans (80, 81). Although mAb PspA16 has limited serotype breadth, it is unclear if other human mAbs to PspA, even those targeting the N-terminal fragment, will be more broadly reactive. It is well established that the N-terminal region of PspA is more variable compared to the proline-rich region, and further studies will determine whether other PspA mAbs are more broadly-reactive than PspA16, as mAb PspA16 binds outside the highly conserved proline-rich region (82). Limitations of this current study include the limited number of mAb isolated for PhtD and PspA. Further isolation of mAbs will identify if the epitopes and gene usage of the mAbs described here are common in multiple donors.

Anti-pneumococcal mAbs have potential for use in the clinic, as current vaccines cover only a subset of current serotypes (although the most prevalent in invasive disease), and a rise in non-vaccine serotypes has occurred following vaccine introduction (24, 83). The prophylactic efficacy of mAbs PhtD3 and PhtD8 were demonstrated against pneumococcal serotype 3, a leading cause of invasive pneumococcal disease (24). We have also assessed the prophylactic efficacy of mAb PhtD3 against serotype 4 in both intranasal and intravenous infection studies, to model pneumococcal pneumonia and sepsis. Furthermore, we have demonstrated the mAb PhtD3 prolongs survival of mice treated with the mAb 24 hrs after infection with pneumococcal serotype 3. As mAbs PhtD3 and PhtD8 target unique epitopes on the N-terminal and C-terminal region of PhtD, respectively, the higher survival of mAb PhtD3-treated mice suggests a potential role of epitope specificity in protective efficacy, although other factors such as functional activity, decreased binding to serotype 3 bacteria in the ELISA and flow binding assays compared to mAb PhtD3, CDR length and percent somatic hypermutation, and corresponding binding modes may be important for the observed differences. Further studies will need to be completed to determine the efficacy of other PhtD mAbs, to examine whether the specific epitope on PhtD indeed influences the protective efficacy of these mAbs, and to determine whether the mAbs protect against infection with additional serotypes. In addition, the delivery timing of the mAbs for optimal protection, the potential use of mAbs in combinations for improved protective efficacy, and the use of mAbs in concert with antibiotic treatment will need to be examined. Furthermore, as secondary pneumococcal infection is prominent following influenza (10) and SARS-CoV-2 infection (11), another potentially useful scenario for use of anti-pneumococcal mAbs would be administration following primary viral infection, to prevent secondary pneumococcal infection. Further studies will need to be completed to determine if mAb PhtD3 or other mAbs will protect against secondary infection.

A potential mechanism of protection for mAbs PhtD3 and PhtD8, and the functional activity of the other PhtD- and PspA-specific mAbs were assessed in opsonophagocytic assays. While showing activity in these *in vitro* assays, the mechanism of protection *in vivo* was not determined and will need to be further explored. mAbs to the pneumococcal capsular polysaccharide have been shown to be protective through multiple mechanisms, with even non-opsonic mAbs demonstrating protective efficacy and the ability to reduce pneumococcal colonization (84, 85). PhtD has been shown to be important for pneumococcal adherence (36, 86), and mAbs to PhtD have also been shown to limit adherence of bacteria (47). Therefore, additional protective mechanisms for anti-PhtD mAbs exist and may work in concert with opsonophagocytosis.

Overall, our study furthers the premise of using human mAbs to highly conserved surface antigens for prevention and treatment of pneumococcal infection. In addition, the application of human mAbs for other bacterial infections, particularly those that have concerns of antibiotic resistance is an important path forward for the development of new therapies. Further defining the epitope specificity of protective human mAbs to conserved pneumococcal surface proteins would also facilitate the development of an epitope-based, and potentially multi-antigen and broadly protective pneumococcal vaccine.

## Material and methods

### Ethics statement

This study was approved by the University of Georgia Institutional Review Board as STUDY00005127 and STUDY00005368. Healthy human donors were recruited to the University of Georgia Clinical and Translational Research Unit and written informed consent was obtained. For the Prevnar-13 vaccinated human samples, healthy subjects were recruited for vaccination with Prevnar-13, and a single blood sample was collected 21-28 days following immunization. All animal studies performed were in accordance with protocols approved by the Institutional Animal Care and Use Committee of the University of Georgia.

### Blood draws and isolation of PBMCs

After obtaining informed consent, 90 mL of blood was drawn by venipuncture into 9 heparin-coated tubes, and 10 mL of blood was collected into a serum separator tube. Peripheral blood mononuclear cells (PBMCs) were isolated from human donor blood samples using Ficoll-Histopaque density gradient centrifugation, and PBMCs were frozen in the liquid nitrogen vapor phase until further use.

### Pneumococcal protein cloning and expression

PspA and PhtD full-length proteins and fragments were cloned from the genome of *S. pneumoniae* strain TCH8431 (serotype 19A) with primers listed in **Table 2** below. The full-length PspA and PhtD were ligated into the pET28a vector while the fragments were ligated into the pMAL-c5x vector. The sequences of all constructed plasmids were confirmed by sequencing, and then transformed into *E. coli* BL21(DE3) for protein expression. Single colonies of transformed *E. coli* were picked and cultured in 5 mL of LB medium supplemented with antibiotic (50 μg/ml kanamycin for pET28a, 100 μg/ml ampicillin for pMAL-c5x) overnight in a shaking incubator at 37 °C. The overnight culture was then expanded at a 1:100 ratio in 2x YT medium with antibiotic and cultured at 37 °C. After the OD_600_ reached 0.5 to 0.7, the culture was induced with 50 μM isopropyl-D-thiogalactopyranoside for 12-16 hrs at room temperature.

**Table 2:**
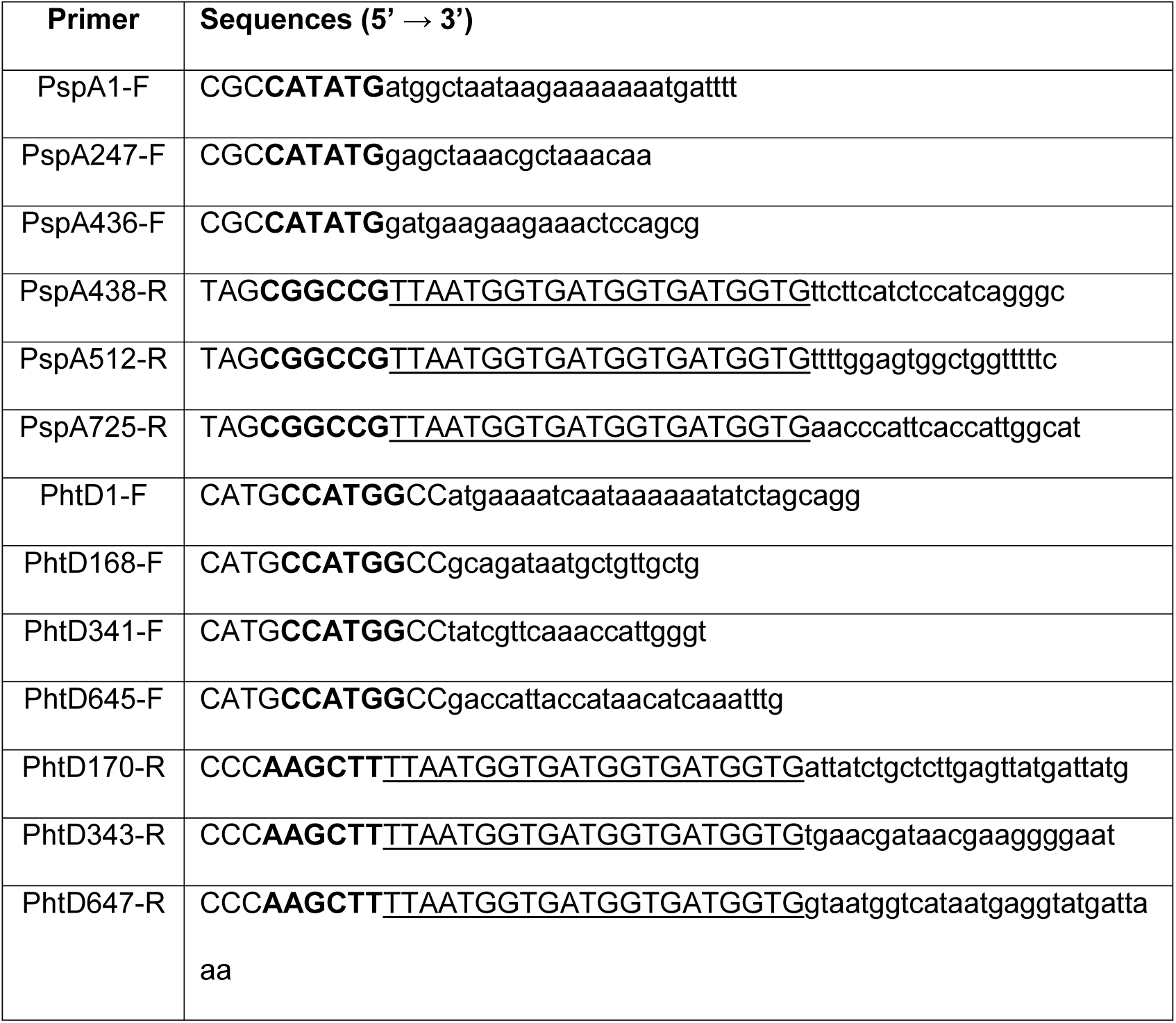
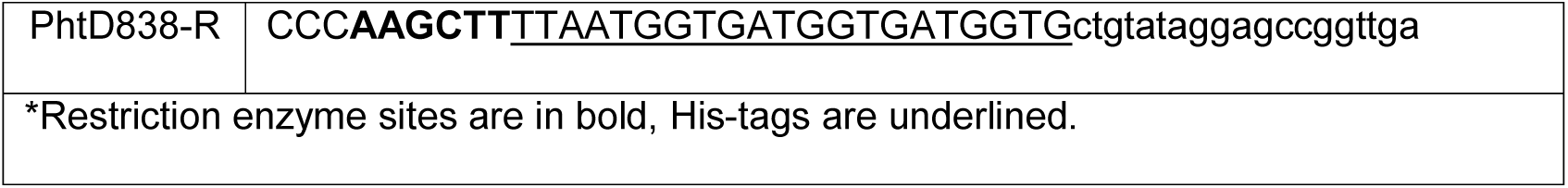
Summary of primers used for cloning of PspA and PhtD genes.

Bacteria pellets collected by centrifugation at 6,000 *x* g for 10 min, and frozen at -80 °C. Thawed *E. coli* pellets were resuspended in 10 mL of buffer containing 20 mM Tris pH 7.5 and 500 mM NaCl, and then lysed by sonication. Cell lysates were centrifuged at 12,000 x g for 30 min and the supernatant was subsequently used for protein purification through a HisTrap column (His- tagged full-length proteins, GE Healthcare) or Amylose resin (MBP-tagged fragments, New England Biolabs) following the manufacturer’s protocols.

### Enzyme-linked immunosorbent assay for binding to pneumococcal proteins

For recombinant protein capture ELISA assays, 384-well plates were treated with 2 µg/ml of antigen in PBS for 1 hr at 37 °C or overnight at 4 °C. Following this, plates were washed once with distilled water before blocking for 1 hr with 2% nonfat milk/2% goat serum in 0.05% PBS-T (blocking buffer). Plates were washed with water three times before applying serially diluted hr. Following this, plates were washed with water three times before applying 25 μL of secondary antibody (goat anti-human IgG Fc; Meridian Life Science) at a dilution of 1:4,000 in blocking solution. After incubation for 1 hr, the plates were washed five times with PBS-T, and 25 μL of a PNPP (p-nitrophenyl phosphate) solution (1 mg/ml PNPP in 1 M Tris base) was added to each well. The plates were incubated at room temperature for 1 hr before reading the optical density at 405 nm on a BioTek plate reader. Binding assay data were analyzed in GraphPad Prism using a nonlinear regression curve fit and the log(agonist)-versus-response function to calculate the binding EC_50_ values.

### Generation of pneumococcal-specific hybridomas

For hybridoma generation, 10 million peripheral blood mononuclear cells purified from the blood of human donors were mixed with 8 million previously frozen and gamma irradiated NIH 3T3 cells modified to express human CD40L, human interleukin-21 (IL-21), and human BAFF(71) in 80 mL StemCell medium A (StemCell Technologies) containing 6.3 µg/mL of CpG (phosphorothioate-modified oligodeoxynucleotide ZOEZOEZZZZZOEEZOEZZZT, Invitrogen) and 1 µg/mL of cyclosporine A (Millipore-Sigma). The mixture of cells was plated in four 96-well plates at 200 µl per well in StemCell medium A. After 6 days, culture supernatants were screened by ELISA for binding to recombinant pneumococcal protein, and cells from positive wells were electrofused to generate hybridomas and biologically cloned as previously described (71).

### Human mAb expression and purification

For hybridoma-derived mAbs, hybridoma cell lines were expanded in StemCell medium A until 80% confluent in 75-cm^2^ flasks. Cells from one 75-cm^2^ cell culture flask were collected with a cell scraper and expanded to 225-cm^2^ cell culture flasks in serum-free medium (Hybridoma-SFM; Thermo Fisher Scientific). Recombinant cultures from transfection were stopped after 5 to 7 days, and hybridoma cultures were stopped after 30 days. For recombinant PhtD3-IgG_2a_, plasmids encoding cDNAs for the heavy and light chain sequences of PhtD3-IgG_2a_ were synthesized (GenScript), and cloned into pCDNA3.1+. mAbs were obtained by transfection of plasmids into Expi293F cells by transfection. For each milliliter of transfection, 1 μg of 25,000-molecular-weight polyethylenimine (PEI; PolySciences Inc.) in 66.67 μL Opti-MEM cell culture medium (Gibco). After 30 min, the DNA-PEI mixture was added to the Expi293F cells, and cells were cultured for 5-6 days for protein expression. Culture supernatants from both approaches were filtered using 0.45 µm filters to remove cell debris. mAbs were purified directly from culture supernatants using HiTrap protein G columns (GE Healthcare Life Sciences) according to the manufacturer’s protocol.

### Isotype determination for human mAbs

For determination of mAb isotypes, 96-well Immulon 4HBX plates (Thermo Fisher Scientific) were coated with 2 µg/mL of each mAb in PBS (duplicate wells for each sample). The plates were incubated at 4 °C overnight and then washed once with water. Plates were blocked with blocking buffer and then incubated for 1 hr at room temperature. After incubation, the plates were washed three times with water. Isotype-specific antibodies obtained from Southern Biotech (goat anti-human kappa-alkaline phosphatase [AP] [catalog number 100244-340], goat anti-human lambda-AP [catalog number 100244-376], mouse anti-human IgG1 [Fc]-AP [catalog number 100245714], mouse anti-human IgG2 [Fc]-AP [catalog number 100245-734], mouse anti-human IgG3 [hinge]-AP [catalog number 100245-824], and mouse anti-human IgG4 [Fc]- AP [catalog number 100245-812]) were diluted 1:1,000 in blocking buffer, and 50 µl of each solution was added to the respective wells. Plates were incubated for 1 h at room temperature and then washed five times with PBS-T. The PNPP substrate was prepared at 1 mg/mL in substrate buffer (1 M Tris base, 0.5 mM MgCl_2_, pH 9.8), and 100 µl of this solution was added to each well. Plates were incubated for 1 hr at room temperature and read at 405 nm on a BioTek plate reader.

### RT-PCR for hybridoma mAb variable gamma chain and variable light chain

RNA was isolated from expanded hybridoma cells using the ENZA total RNA kit (Omega BioTek) according to the manufacturer’s protocol. cDNA was obtained using the Superscript IV Reverse Transcriptase kit. Following this, PCR was conducted in two steps using established primers for the heavy chain, and kappa and lambda light chains (87). Samples were analyzed by agarose gel electrophoresis and purified PCR products (ENZA cycle pure kit; Omega Bio- Tek) were cloned into the pCR2.1 vector using the Original TA cloning kit (Thermo Fisher Scientific) according to the manufacturer’s protocol. Plasmids were purified from positive DH5α colonies with ENZA plasmid DNA mini kit (Omega Bio-Tek) and submitted to Genewiz for sequencing. Sequences were analyzed using IMGT/V-Quest (88).

### Experimental setup for biolayer interferometry

For all biosensors, an initial baseline in running buffer (PBS, 0.5% bovine serum albumin [BSA], 0.05% Tween 20, 0.04% thimerosal) was obtained. For epitope mapping, 100 µg/mL of His-tagged PhtD protein was immobilized on anti-penta-HIS biosensor tips (FortéBio) for 120 s. For binding competition, the baseline signal was measured again for 60 s before biosensor tips were immersed into wells containing 100 µg/mL of primary antibody for 300 s. Following this, biosensors were immersed into wells containing 100 µg/mL of a second mAb for 300 s. Percent binding of the second mAb in the presence of the first mAb was determined by comparing the maximal signal of the second mAb after the first mAb was added to the maximum signal of the second mAb alone. mAbs were considered noncompeting if maximum binding of the second 66% of its uncompeted binding. A level of between ≥33% and 66% of its uncompeted binding was considered intermediate competition, and ≤

### Bacterial strains and growth conditions

Pneumococcal strains were grown at 37 °C in 5% CO_2_ in Todd-Hewitt broth (BD, Franklin Lakes NJ) supplemented with 0.5% yeast extract for 12 hrs. Ten percent glycerol was added to the media and 500 µL aliquots were made. Cultures were kept at -80 °C until used, cultures were washed twice with PBS before being used in experiments. Colonies were grown on BD Trypticase Soy Agar II with 5% Sheep Blood (BD, Franklin Lakes NJ). The numbers of CFUs per milliliter of these stocks were determined, after the aliquots had been frozen, by plating a single quick-thawed diluted aliquot on sheep’s blood agar plates. The calculated number of CFUs was subsequently used to make dilutions for experiments from aliquots thawed at later times. In each experiment, the actual number of CFUs administered was determined by plating on blood agar at the time of the assay. Strains used in this study are listed in **Table 3**.

**Table 3:**
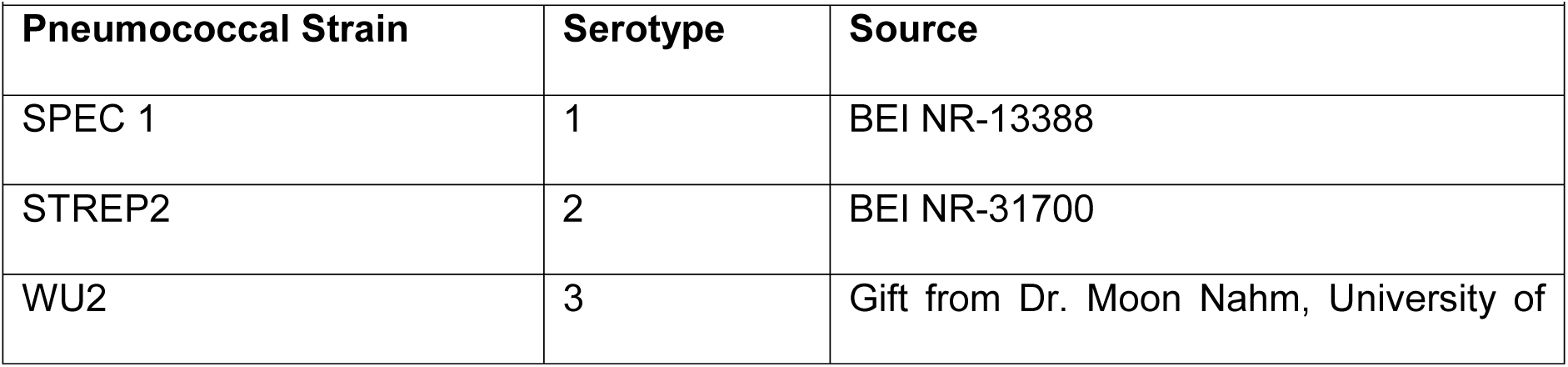
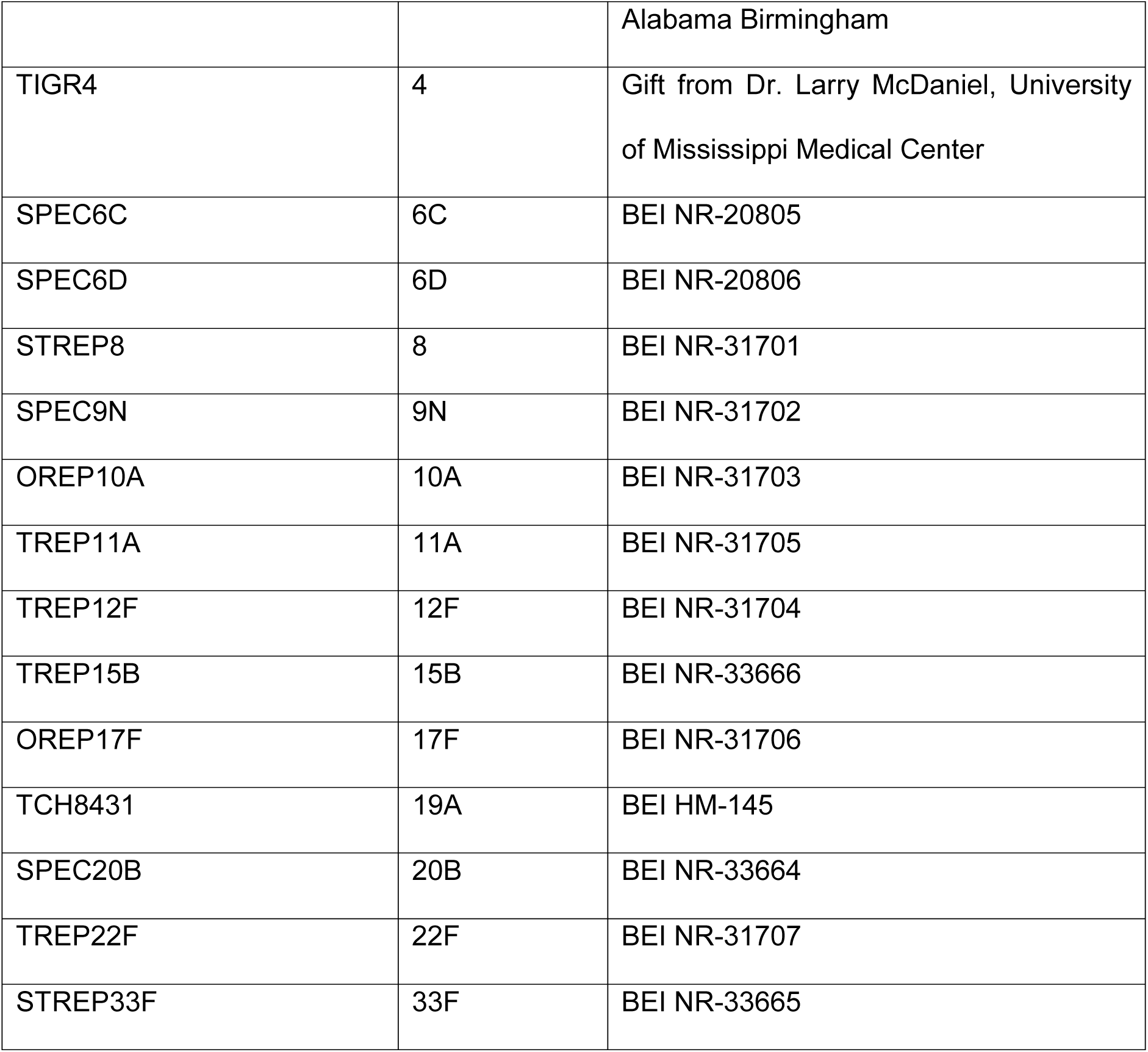
Summary of pneumococcal strains used in this study.

### Western blot

Pneumococcal strains were mixed with non-reducing loading buffer (Laemmli SDS sample buffer, non-reducing 6X) and loaded on a 4-12% Bis-Tris gel (Invitrogen). Samples were then transferred to PVDF membranes via iBlot system (Invitrogen) and then blocked with 5% blocking buffer (5% nonfat milk in PBS-T) for 1 hr at room temperature or at 4°C overnight. The membrane was washed three times in five-minute intervals on an orbital shaker with 0.05% PBS-T. Then, primary antibodies were added at dilutions of 1 µg/mL in PBS for one hour at room temperature. The membranes were then washed three time in five-minute intervals with PBS-T on an orbital shaker, and soaked in the secondary antibody at a 1:8,000 dilution in blocking buffer for one hour. Next, the membranes were then washed five times in five-minute intervals on the orbital shaker with PBS-T, and substrate (Pierce ECL Western Blotting Substrate, Thermo Scientific) was added and an image was taken immediately with the ChemiDoc Imaging System (BioRad).

### Enzyme-linked immunosorbent assay of fixed pneumococcus

384-well plates were treated with 15 µL (∼10^7^ CFUs) of whole cell pneumococcus in PBS into each well. Cell density was checked by microscope to ensure a confluent layer of pneumococcus was coated. The bacteria were then fixed with 15 µl of 4% paraformaldehyde into each well and placed onto a plate shaker for 10 mins to mix. The 384-well plates were incubated at 4 °C for 24-48 hours to allow the bacteria to fix to the bottom of the plates. Following this, the plates were washed once with 75 µl of PBS-T into each well. The plates were then blocked with 2% blocking buffer for 1 hr at room temperature then washed three times with PBS-T. Next, 25 µl of serially diluted primary antibodies were applied to the wells for 1 hr at room temperature, then plates were washed with PBS-T three times. Following this, 25 µl of secondary antibody (goat anti-human IgG Fc; Meridian Life Science) at a 1:4,000 dilution in blocking buffer was applied to each well for 1 hr at room temperature. After the plates were washed with PBS-T five times, 25 µl of PNPP (p-nitrophenyl phosphate) solution (1 mg/ml PNPP in 1 M Tris bases) was added to each well for 1 hr at room temperature. After 1 hr the optical density was read at 405 nm on a BioTek plate reader. Binding assay data were analyzed in GraphPad Prism.

### Binding of antibodies to bacteria by flow cytometry

The ability of mAbs to bind antigen exposed on the surface of *S. pneumoniae* was determined by flow cytometry. Bacteria were stained with 10 µM CFSE (Millipore Sigma) for 1 hr at 37 °C. Bacteria were then washed with Hank’s Balanced Salt Solution (HBSS) containing 1% bovine serum albumin (BSA) to remove excess stain. Following this, 1x10^6^ bacteria were incubated with 10 μg/ml of antibody for 30 min at 37 °C. Bacteria were then washed twice with HBSS+1% BSA. Antibody binding was detected using an APC Anti-Human IgG Fc (Biolegend) at a 1:100 dilution incubated for 1 hr with the bacteria. Cells were washed with HBSS+1% BSA and fixed in 2% paraformaldehyde (PFA) in PBS prior to analysis on a NovoCyte Quanteon Flow Cytometer.

### Determination of mAb efficacy

For intranasal challenge study with TIGR4, 5-7-week-old CBA/CaHN-Btkxid/J (CBA/N) mice (The Jackson Laboratory, Bar Harbor, ME) were used. Mice were intraperitoneally inoculated with antibody treatments 2 hrs prior to pneumococcal infection. For infection, mice were anesthetized by inhalation of 5% isoflurane and intranasally challenged with 40 µL of PBS containing 10^5^ colony-forming units (CFUs) of TIGR4. Mice were weighed and assessed daily, and were considered moribund when >20% of body weight was lost or they were nonresponsive to manual stimulation or exhibited respiratory distress. Mice were euthanized by CO_2_ asphyxiation followed by cervical dislocation. For the intravenous challenge with TIGR4, C57BL/6 mice 5-7 weeks old (Charles River) were used. Mice were intraperitoneally inoculated with antibody treatments two hrs prior to pneumococcal infection, and infected intravenously with 10^6^ CFUs of TIGR4 via the tail vein. Mice were monitored and euthanized as described above. For intranasal challenge studies with WU2, C57BL/6 mice 5-7 weeks old (Charles River) were used. For intranasal infection, mice were anesthetized by inhalation of 5% isoflurane and intranasally challenged with 40 µL of PBS containing 10^6^ colony-forming units (CFUs) of WU2. Mice were either treated 2 hrs before infection or 24 hours post infection by intraperitoneally inoculating with antibody. In prophylactic studies, mice were euthanized based on the humane endpoints above. For treatment studies, mice were euthanized when >30% of pre-infection body weight was lost or they were nonresponsive to manual stimulation or exhibited respiratory distress. Actual doses delivered to mice in all studies were determined by titering the bacteria after delivery.

### Opsonophagocytic killing assay

An opsonophagocytic killing assay was performed as described previously (89, 90) as adapted from an earlier protocol with modifications (91). TIGR4 stocks were incubated in triplicate wells in a 96-well round-bottom plate for 1 hour at 37°C with the indicated antibodies (10 µg of antibody per well in a final volume of 100 µL per well) in opsonization buffer B (OBB: sterile 1× PBS with Ca^2+^/Mg^2+^, 0.1% gelatin, and 5% heat-inactivated FetalClone [HyClone]), with heat-inactivated FetalClone-treated only TGR4 cells serving as a control. Cells of the human promyelocytic leukemia cell line HL-60 (ATCC) were cultured in RPMI with 10% heat-inactivated FetalClone and 1% l-glutamine. HL-60 cells were differentiated using 0.6% *N*,*N*- dimethylformamide (DMF [Fisher]) for 3 days before performing the OPA assay, harvested, and resuspended in OBB. Baby rabbit complement (Pel-Freez) was added to HL-60 cells at a 1:5 final volume. The HL-60–complement mixture was added to the bacteria at 5 × 10^5^ cells/well. The final reaction mixtures were incubated at 37**°**C for 1 hour with shaking. The reactions were stopped by incubating the samples on ice for approximately 20 min. Then 10 μ of each reaction mixture (triplicate) was diluted to a final volume of 50 μL and plated onto blood agar plates. Plates were incubated overnight at 30**°**C and counted the next day. The percentage of bacterial killing was calculated as each sample replicate normalized to the mean value obtained for the control samples, subtracted from 100 (with No Ab control samples representing 0% survival).

### Flow-based opsonophagocytosis assay

Pneumococcal cells were stained with pHRodo Succinimidyl Ester (Invitrogen) following manufacturer’s protocol. Approximately, ∼10^8^ CFUs of bacteria were fixed with 1% paraformaldehyde in PBS for 30 min at room temperature. Fixed bacteria were washed twice with PBS and resuspended with 0.5 mL freshly prepared 100 mM NaHCO_3_ (pH 8.5). Immediately before use, the contents of a 0.1 mg vial of pHRodo iFL amine-reactive dye were dissolved in 10 μL of DMSO to prepare a 10 mM stock solution. pHRodo was diluted in the bacterial suspension at a final concentration of 0.1 mM, and bacteria were stained for 1 hr at room temperature. Stained bacteria were washed twice with Hank’s Balanced Salt Solution with Ca^2+^ and Mg^2+^ (HBSS, Gibco), and resuspended with 0.5 mL HBSS and stored in the dark at 4 °C. The opsonophagocytosis assay was performed in 96 well U-bottom plates in a total volume of 120 μL per well. First, 20 μL of pHRodo labeled bacteria (∼10^7^ CFUs/well) was mixed with 40 μL of sterile filtered mAbs (50 μg/well), and incubated on a shaker at 37 °C for 30 min. Bacteria were mixed with HBSS as a negative control, and purified human serum IgG was used as a positive control. Differentiated HL-60 cells were washed twice with HBSS and mixed with baby rabbit complement (Pel-Freez Biologicals) at a final concentration of 10% in each well. Following this, 60 μL (1×10^6^ viable cells) of differentiated HL60 cells and complement were added to the mixture of bacteria and antibodies, and incubated on a shaker at 37 °C for 60 min. The plate was then centrifuged at 1300 rpm for 5 min at 4 °C to remove the supernatant and the pellet was washed twice with 200 μL of HBSS. After the second wash, the pellet was resuspended in a 50 μL mixture of PE-anti-human CD11b (Southern Biotech, 10 μL/million cells), Alexa Fluor 647-anti-human CD35 (BD Biosciences, 5 μL/million cells), and DAPI (Invitrogen, 50 ng/million cells) in PBS containing 1% BSA. After a 30 min incubation at 4 °C in the dark, the plate was washed twice with 200 μL of PBS, and cells were resuspended in 100 μL of PBS. Cells were analyzed with a NovoCyte Quanteon Flow Cytometer. Single fluorophore stained differentiated HL60 cells and pHRodo stained bacteria were used to calculate the compensation matrix. A total of 10,000 ungated events were collected from each sample well, and data were analyzed by FlowJo.

## Supporting information

Supplemental antibody sequences

## Acknowledgements

These studies were supported by startup funding from the University of Georgia Office of the Vice President for Research, a Junior Faculty Seed Grant from the University of Georgia Office of Research, and National Institutes of Health grants 1K01OD026569 (JJM) and R01AI123383 (FYA). Human subject studies were partially funded by the National Center for Advancing Translational Sciences award number UL1TR002378. FR was supported by National Institutes of Health NIGMS grant GM109435, Post-Baccalaureate Training in Infectious Diseases Research. The funders had no role in study design, data collection and analysis, decision to publish, or preparation of the manuscript. The content is solely the responsibility of the authors and does not necessarily represent the official views of the National Institutes of Health. We thank the University of Georgia Clinical and Translational Research Unit for assistance with donor identification and blood draws, and the University of Georgia Center for Tropical and Emerging Global Diseases and College of Veterinary Medicine flow cytometry cores for assistance with cell sorting and analysis.

