## Supplemental antibody sequences for "Broadly reactive human monoclonal antibodies targeting the pneumococcal histidine triad protein protect against fatal pneumococcal infection"

**mAb PhtD3**

Heavy chain variable region nucleotide sequence:

caggtgcagctagtgcagtctgggcctgacgtgaagaagcctgggtcctcggtgaaggtctcctgcaaggcctctggagccgccttcgagagttttgccttcgcctgggtgcgacaggcccctggacaagggtttgagtggatgggaaggatcattccaatcttggaaacaagggactacgcagagaagttccagggcagaatgacgatgaccacagacgagtcgacggcgacagcctacatggaactgaacagcctaagatttgaagacacggccgtttatttctgtgcgcgagatgggcacattatgaggacaactctctcggatgctgcacttgatgtctggggccaagggacaacggtcattgtctcctcag

Heavy chain variable region amino acid sequence:

QVQLVQSGPDVKKPGSSVKVSCKASGAAFESFAFAWVRQAPGQGFEWMGRIIPILETRDYAEKFQGRMTMTTDESTATAYMELNSLRFEDTAVYFCARDGHIMRTTLSDAALDVWGQGTTVIVSS

Light chain variable region nucleotide sequence:

gacatcgtgatgacccagtctccagtcaccctgtctttgtctccaggggagagagccaccctctcctgcagggccagtcagagtcttactgacaactacttagcctggtaccagcagaaacctggccaggctcccaggctcctcatctacgccgcatccaccagggccactggcatcccagacaggatcagtggcagtgggtctgggacagacttcactctcaccatcagcagagtggagcctgaagattttgcaatgttttactgtcaacagtatcagaactcaccgttcaccttcggcggggggaccacggtggagatcaaac

Light chain variable region amino acid sequence:

DIVMTQSPVTLSLSPGERATLSCRASQSLTDNYLAWYQQKPGQAPRLLIYAASTRATGIPDRISGSGSGTDFTLTISRVEPEDFAMFYCQQYQNSPFTFGGGTTVEIK

**mAb PhtD6**

Heavy chain variable region nucleotide sequence:

caggtgcagttggtgcagtctgggactgaggtgaagaagcctggggcctcagtgaaggtcgcctgcaaggcttctggatacaccttcactagttatgatatcaactgggtgcgacaggcccctggacaagggcttgagtggatgggatggatgaacgcgaacagtggcaacacaggctatgcacaaaagttccagggcagagtcaccatgaccaggaacacctccattaccacagcctacatggacctgattgatctgacatctgaggacacggccatatattactgtgcgagagggccgtactgggtggagaattggttcgacacctggggccagggaaccctggtcagcgtctcctcag

Heavy chain variable region amino acid sequence:

QVQLVQSGTEVKKPGASVKVACKASGYTFTSYDINWVRQAPGQGLEWMGWMNANSGNTGYAQKFQGRVTMTRNTSITTAYMDLIDLTSEDTAIYYCARGPYWVENWFDTWGQGTLVSVSS

Light chain variable region nucleotide sequence:

gacatccagatgacccagtctccatcctccctgtctgcgtctgtcggagacagagtcaccatcacttgccgggcaagtcggagcattcgcagctttttaaattggtatcaacaaaaaccagggaaaccccctaacctcctgatctataaagcatccactttgcacagtggggtcccgtctaggttcagtggcagtggatctgggacagatttcactctcacaatcaacaatctacaacccgaagattttgcaacttactactgtcaacagagttacagtaatcagaagaccttcggccaagggaccaaggtggacatcaaac

Light chain variable region amino acid sequence:

DIQMTQSPSSLSASVGDRVTITCRASRSIRSFLNWYQQKPGKPPNLLIYKASTLHSGVPSRFSGSGSGTDFTLTINNLQPEDFATYYCQQSYSNQKTFGQGTKVDIK

**mAb PhtD7**

Heavy chain variable region nucleotide sequence:

gaggtgcagctggtgcagtctggggctgaagtgaagaagcctggggcctcagtgaaggtctcctgcaaggcttctggagacatcttcagcgactcctatattcactgggtgcgacaggcccctggacaagggcctgagtggatgggatgggtcagccctaacactggtgccacacattatgcacagaagttgcagggcagagtcaccatgaccagcgacacgtccatcagtacagcctatttggagctgaccaggctggcatctgacgacacggccgtttattactgtgcgagagtcttaaggggaagttatgatttccggggtaattatccacatgattttgactactggggccagggaaccctggtcaccgtctcctcag

Heavy chain variable region amino acid sequence:

EVQLVQSGAEVKKPGASVKVSCKASGDIFSDSYIHWVRQAPGQGPEWMGWVSPNTGATHYAQKLQGRVTMTSDTSISTAYLELTRLASDDTAVYYCARVLRGSYDFRGNYPHDFDYWGQGTLVTVSS

Light chain variable region nucleotide sequence:

cagcttgtgctgactcaaccgccctctgcctctgcctccctgggagcctcggtcaccctcacctgcactctgagcagaggacacaacaactaccccatcgcttggctccaaaagcagacagataagggccctcgttatgtgatgagacttaatagtgatggcagccaccacaagggggacggaatccctgatcgcttctcaggctccagttctggggctgagcgctacctcagcatttccagtctccagcctgaagatgaggctgaatactactgtcagacgtgggacactggccttcagggggtgttcggcggagggaccaaactgttcgtcctag

Light chain variable region amino acid sequence:

QLVLTQPPSASASLGASVTLTCTLSRGHNNYPIAWLQKQTDKGPRYVMRLNSDGSHHKGDGIPDRFSGSSSGAERYLSISSLQPEDEAEYYCQTWDTGLQGVFGGGTKLFVL

**mAb PhtD8**

Heavy chain variable region nucleotide sequence:

caggtgcagctggtgcagtctggggctgaggtgaagaagcctggggcctcagtgaaggtctcctgtaaggcttctggatacaccttcaccgactactttatacactgggtgcgacaggcccctggacacggtcttgaatggatggggtggatcaaccctaaccgcggtgtcacaaactatacacagaagtttcagggcagggtcaccatgaccaaggacacgtccgtcacctcagtctacatggagctgagcaggctgacatctgacgacacggccctatattattgtgcgagaggtggtacgcttgaccactggggccagggcaccctggtcaccgtctcctctg

Heavy chain variable region amino acid sequence:

QVQLVQSGAEVKKPGASVKVSCKASGYTFTDYFIHWVRQAPGHGLEWMGWINPNRGVTNYTQKFQGRVTMTKDTSVTSVYMELSRLTSDDTALYYCARGGTLDHWGQGTLVTVSS

Light chain variable region nucleotide sequence:

cagcttgtgctgactcaatcgccctctgcctctgcctccctgggagcctcggtcaccctcacctgcactctgagcagtgggcacagcacctacgacatcgcatggcatcagcagcagccaggaaagggccctcgacacttgatgagacttaacggtgatggcagtcacaccaacggggacgggatccctgatcgcttctcaggctccagctctggggctgagcgctacctcaccatctccagcctccagtctgaagatgaggctgactattactgtcacacctgggtcactaacattcatttggtgttcggcggagggaccaaactgaccgtcctag

Light chain variable region amino acid sequence:

QLVLTQSPSASASLGASVTLTCTLSSGHSTYDIAWHQQQPGKGPRHLMRLNGDGSHTNGDGIPDRFSGSSSGAERYLTISSLQSEDEADYYCHTWVTNIHLVFGGGTKLTVL

**mAb PspA16**

Heavy chain variable region nucleotide sequence:

caggtgcagctggtgcagtctgggcctgacgtgaagaagcctggggcctcagtgaaggtctcctgtaagacttctggatacaccttcactggctactatatgcactgggtgcgacaggcccctggacaagggcttgagtggatgggatgggtcaaccctaacaccggtggcacaagttatgcacagaagtttcagggcagggtcaccgtgaccagggacacgtccatcagcacagtctacatggaactgagcgctctaggatctgacgacacggccatatatttctgtgcgagggcgtgggctccgggcgctgagtacctccaccactggggccagggcaccctggtcaccgtctcctcag

Heavy chain variable region amino acid sequence:

QVQLVQSGPDVKKPGASVKVSCKTSGYTFTGYYMHWVRQAPGQGLEWMGWVNPNTGGTSYAQKFQGRVTVTRDTSISTVYMELSALGSDDTAIYFCARAWAPGAEYLHHWGQGTLVTVSS

Light chain variable region nucleotide sequence:

gagattgtgatgacccagtctccaggcaccctgtctttgtctccaggggaaacagccaccctctcctgcagggccagtcagagtgttggcagcagctacttagcctggtatcagcagaaacctggccaggctcccaggctcctcatctttggtgcgtccaacagggccactggcatcccagtcaggttcagtgccagtgggtctgggacagacttcactctcaccatcagcagactggagcctgaagatttcgcagtgtattactgtcagcagcatgatcactcaccattcactttcggccctgggaccaaagtggatctcaaac

EIVMTQSPGTLSLSPGETATLSCRASQSVGSSYLAWYQQKPGQAPRLLIFGASNRATGIPVRFSASGSGTDFTLTISRLEPEDFAVYYCQQHDHSPFTFGPGTKVDLK

**PhtD3-IgG_2a_ human/mouse chimera**

Heavy chain amino acid sequence

QVQLVQSGPDVKKPGSSVKVSCKASGAAFESFAFAWVRQAPGQGFEWMGRIIPILETRDYAEKFQGRMTMTTDESTATAYMELNSLRFEDTAVYFCARDGHIMRTTLSDAALDVWGQGTTVIVSSGCTAGCTTAPSVYPLAPVCGDTTGSSVTLGCLVKGYFPEPVTLTWNSGSLSSGVHTFPAVLQSDLYTLSSSVTVTSSTWPSQSITCNVAHPASSTKVDKKIEPRGPTIKPCPPCKCPAPNLLGGPSVFIFPPKIKDVLMISLSPIVTCVVVDVSEDDPDVQISWFVNNVEVHTAQTQTHREDYNSTLRVVSALPIQHQDWMSGKEFKCKVNNKDLPAPIERTISKPKGSVRAPQVYVLPPPEEEMTKKQVTLTCMVTDFMPEDIYVEWTNNGKTELNYKNTEPVLDSDGSYFMYSKLRVEKKNWVERNSYSCSVVHEGLHNHHTTKSFSRTPGK

Light chain amino acid sequence

DIVMTQSPVTLSLSPGERATLSCRASQSLTDNYLAWYQQKPGQAPRLLIYAASTRATGIPDRISGSGSGTDFTLTISRVEPEDFAMFYCQQYQNSPFTFGGGTTVEIKGGATCCRADAAPTVSIFPPSSEQLTSGGASVVCFLNNFYPKDINVKWKIDGSERQNGVLNSWTDQDSKDSTYSMSSTLTLTKDEYERHNSYTCEATHKTSTSPIVKSFNRNEC

**PhtD8-IgG_2a_ human/mouse chimera**

Heavy chain amino acid sequence

QVQLVQSGAEVKKPGASVKVSCKASGYTFTDYFIHWVRQAPGHGLEWMGWINPNRGVTNYTQKFQGRVTMTKDTSVTSVYMELSRLTSDDTALYYCARGGTLDHWGQGTLVTVSSASTTAPSVYPLAPVCGDTTGSSVTLGCLVKGYFPEPVTLTWNSGSLSSGVHTFPAVLQSDLYTLSSSVTVTSSTWPSQSITCNVAHPASSTKVDKKIEPRGPTIKPCPPCKCPAPNLLGGPSVFIFPPKIKDVLMISLSPIVTCVVVDVSEDDPDVQISWFVNNVEVHTAQTQTHREDYNSTLRVVSALPIQHQDWMSGKEFKCKVNNKDLPAPIERTISKPKGSVRAPQVYVLPPPEEEMTKKQVTLTCMVTDFMPEDIYVEWTNNGKTELNYKNTEPVLDSDGSYFMYSKLRVEKKNWVERNSYSCSVVHEGLHNHHTTKSFSRTPGK

Light chain amino acid sequence

QLVLTQSPSASASLGASVTLTCTLSSGHSTYDIAWHQQQPGKGPRHLMRLNGDGSHTNGDGIPDRFSGSSSGAERYLTISSLQSEDEADYYCHTWVTNIHLVFGGGTKLTVLGSPKSTPTLTVFPPSSEELKENKATLVCLISNFSPSGVTVAWKANGTPITQGVDTSNPTKEGNKFMASSFLHLTSDQWRSHNSFTCQVTHEGDTVEKSLSPAECL
